# Single cell regulatory architecture of human pancreatic islets suggests sex differences in β cell function and the pathogenesis of type 2 diabetes

**DOI:** 10.1101/2024.04.11.589096

**Authors:** Mirza Muhammad Fahd Qadir, Ruth M. Elgamal, Keijing Song, Parul Kudtarkar, Siva S.V.P Sakamuri, Prasad V. Katakam, Samir El-Dahr, Jay Kolls, Kyle J. Gaulton, Franck Mauvais-Jarvis

## Abstract

Biological sex affects the pathogenesis of type 2 and type 1 diabetes (T2D, T1D) including the development of β cell failure observed more often in males. The mechanisms that drive sex differences in β cell failure is unknown. Studying sex differences in islet regulation and function represent a unique avenue to understand the sex-specific heterogeneity in β cell failure in diabetes. Here, we examined sex and race differences in human pancreatic islets from up to 52 donors with and without T2D (including 37 donors from the Human Pancreas Analysis Program [HPAP] dataset) using an orthogonal series of experiments including single cell RNA-seq (scRNA-seq), single nucleus assay for transposase-accessible chromatin sequencing (snATAC-seq), dynamic hormone secretion, and bioenergetics. In cultured islets from nondiabetic (ND) donors, in the absence of the *in vivo* hormonal environment, sex differences in islet cell type gene accessibility and expression predominantly involved sex chromosomes. Of particular interest were sex differences in the X-linked KDM6A and Y-linked KDM5D chromatin remodelers in female and male islet cells respectively. Islets from T2D donors exhibited similar sex differences in differentially expressed genes (DEGs) from sex chromosomes. However, in contrast to islets from ND donors, islets from T2D donors exhibited major sex differences in DEGs from autosomes. Comparing β cells from T2D and ND donors revealed that females had more DEGs from autosomes compared to male β cells. Gene set enrichment analysis of female β cell DEGs showed a suppression of oxidative phosphorylation and electron transport chain pathways, while male β cell had suppressed insulin secretion pathways. Thus, although sex-specific differences in gene accessibility and expression of cultured ND human islets predominantly affect sex chromosome genes, major differences in autosomal gene expression between sexes appear during the transition to T2D and which highlight mitochondrial failure in female β cells.

## Introduction

Type 1 and type 2 diabetes (T1D, T2D) are heterogeneous diseases and biological sex affects their pathogenesis. In the context of T2D, sex affects the development of adiposity, insulin resistance, and dysfunction of insulin-producing β cells of pancreatic islets.^1,2^ For example, ketosis-prone diabetes is a form of T2D with acute β cell failure and severe insulin deficiency predominantly observed in black men.^3–5^ A missense mutation in the β cell-enriched MAFA transcription factor is found in subjects with adult-onset β cell dysfunction, where men are more prone to β cell failure than women.^6^ Similarly, T1D is the only common autoimmune disease characterized by a male predominance^1,7–9^, and males who develop T1D during puberty have lower residual β cell function than females at diagnosis.^10^ Furthermore, among T1D subjects receiving pancreatic islet transplantation, recipients of male islets exhibit early graft β cell failure when compared to recipients of female islets.^11^ The mechanisms that drive preferential β cell failure in males, however, is unknown. Studying sex differences in islet biology and dysfunction represent a unique avenue to understand sex-specific heterogeneity in β cell failure in diabetes.^2^

Female- and male-specific blood concentrations of the gonadal hormones estradiol and testosterone produce differences in islet function *in vivo*.^12–20^ However, the sex-specific and cell autonomous factors that influence islet function outside the *in vivo* hormonal environment are unknown. These differences could be due to sex chromosome gene dosage, or epigenetic programming caused by testicular testosterone during development in males.^1,21,22^ The Genotype-Tissue Expression (GTEx) project analysis of the human transcriptome across various tissues revealed that the strongest sex bias is observed for X-chromosome genes showing higher expression in females.^23^ In the pancreas, the majority of genes with sex-biased expression are on the sex chromosomes and most sex-biased autosomal genes are not under direct influence of sex hormones.^24^ In human pancreatic islets, DNA methylation of the X-chromosome is higher in female than males.^25^ Thus, the cell autonomous influence of sex chromosome genes may impact sex-specific islet biology and dysfunction and diabetes pathogenesis.

Here, we examined sex and race differences in human pancreatic islets from up to 52 donors with and without T2D using an orthogonal series of experiments including single cell RNA-seq (scRNA-seq), single nucleus assay for transposase-accessible chromatin sequencing (snATAC-seq), and dynamic hormone secretion and bioenergetics. Our studies establish biological sex as a genetic modifier to consider when designing experiments of islet biology.

## Results

### Human islet cells show conserved autosomal gene expression signatures independent of sex and race

We performed scRNA-seq on pancreatic islets from age- and BMI-matched non-diabetic donors across race and sex (Tulane University Islet Dataset, TUID, n=15), which we combined with age- and BMI-matched non-diabetic donors and donors with T2D from the HPAP database^26,27^ (n=37) to create an integrated atlas of islet cells **(Fig. 1a and Extended Data Fig. 1a-b)**. To obtain high-quality single cell signatures, we used a series of thresholds including filtering, ambient RNA correction, and doublet removal, resulting in 141,739 high-quality single cell transcriptomes, with TUID showing optimal sequencing metrics **(Extended Data Fig. 1c and 1d)**. We identified 17 cell clusters, which we annotated based on marker genes with differential expression (DEGs) correlating to known transcriptional signatures of islet cells **(Fig. 1b)**.^28^ Cell clusters showed even distribution across sex, race, disease, and library of origin **(Fig. 1c)**. Consistent with a prior analysis^26^, all islet cell clusters except for lymphocytes and Schwann cells were identified in HPAP data **(Extended Data Fig. 1b)**. Notably, we observed greater variability in total cell number within each donor library in HPAP compared to TUID **(Fig. 1d)**. We observed a high degree of correlation between cell-specific gene expression and cell clusters across donors **(Extended Data Fig. 1e)**. As expected, sex chromosome-specific transcripts were expressed across male and female cell types **(Extended Data Fig. 1f)**.

**Fig. 1:**
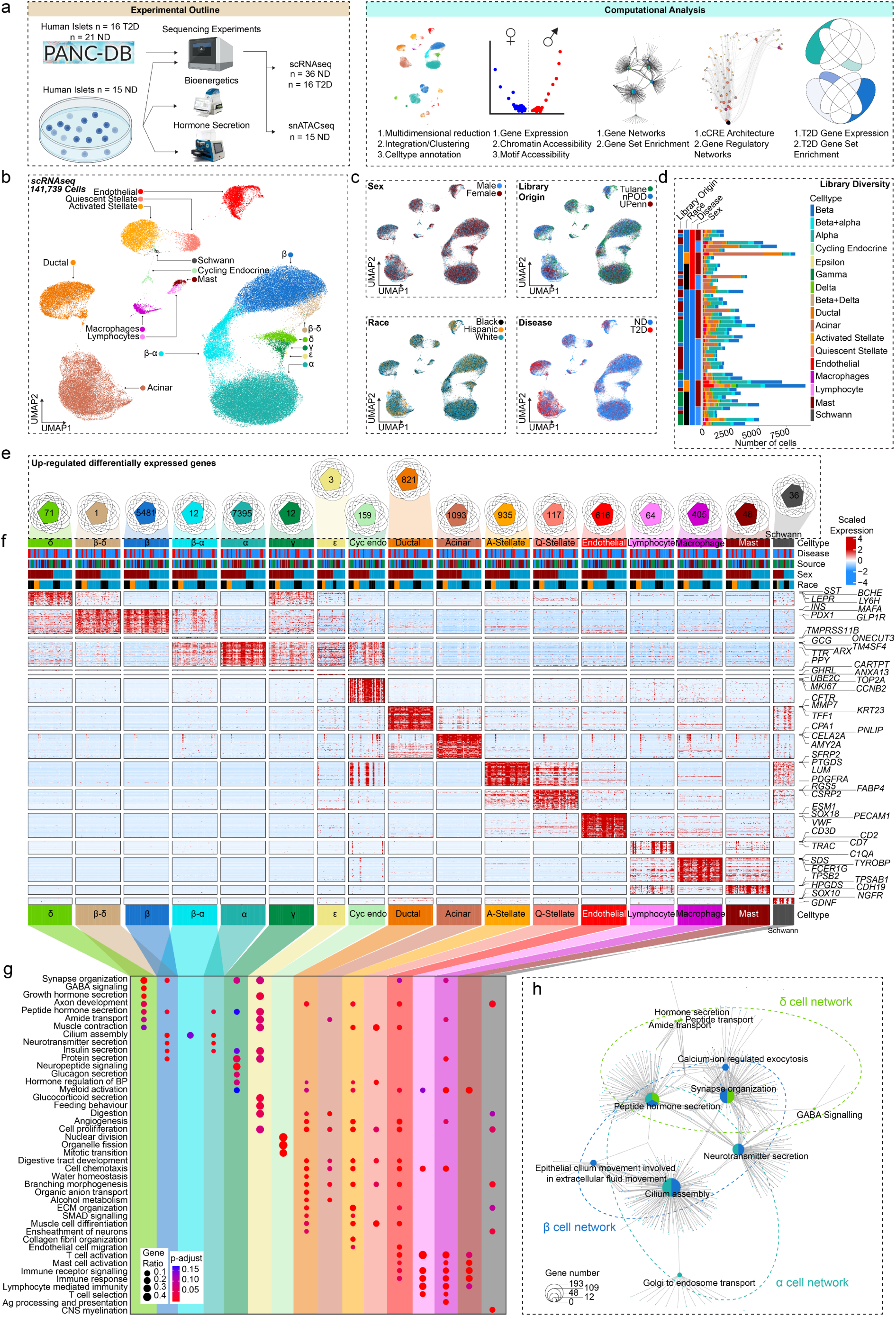
Pancreatic islet cells have a conserved expression signature across sex and race. **a,** Experimental and computational design. **b,** UMAP plot denoting integrated clustering of 141,739 single pancreatic islet cells across 17 clustered cell types based on their scRNAseq profiles, spanning n=52 datasets. Each cluster cell type is denoted by a label and color. **c,** Cells diversified based on donor’s sex, origin, race, and disease status. **d,** Cell number stemming from each of the n=52 donors, grouped based on origin, race, disease status and sex. **e,** Venn diagrams showing conserved differentially expressed genes (DEGs) upregulated in each cluster, across race and sex in non-diabetic donors. Each number denotes conserved upregulated genes across sex and race. Venn diagram identities are colored based on clusters shown in A. **f,** Gene expression heatmap of conserved genes grouped based on colored and labelled clusters as in A. Heatmap is grouped based on disease, source, sex, and race, as denoted by the bars on top. Select genes are labeled on the y-axis. **g,** Gene ontology (GO) analysis showing select upregulated pathways across clusters as shown in E. The intensity of the color denotes scaled FDR corrected adj p-value, and size of the bubble denotes the gene:query ratio. **h,** Activated pathway network analysis of conserved pathways across sex and race in case of β, α and δ cell clusters. n= 36 non-diabetic and n=16 T2D diabetic donors. DEGs have FDR adjusted q-value<0.1, GO pathways have FDR adjusted q-value<0.2

We more broadly examined DEGs across clusters by creating sample ‘pseudo-bulk’ profiles for each cell type to control for pseudo-replication of cells being repetitively sampled from a fixed donor. For example, each β cell per donor was aggregated into one profile, enabling us to control for the disproportionate β cell numbers across donors **(Fig. 1d)**. Autosomal genes with expression specific to each cell cluster were consistent across sex and race. In endocrine cell types, we found 5,481 β (*INS*, *MAFA*), 7,395 α (*GCG*, *ARX*), 71 δ (*HHEX*, *SST*), 3 ε (*GHRL*), 12 γ (*PPY*) and 159 cycling endocrine (*TOP2A*, *MKI67*) DEGs **(Fig. 1e-f and Extended Data Fig. 1g)**. In non-endocrine cell types, we found 821 ductal (*CFTR*, *TFF1*), 1,093 Acinar (*PNLIP*, *AMY2A*), 117 quiescent stellate (*PTGDS*, *DCN*), 935 activated stellate (*RGS5*, *FABP4*), 616 endothelial (*PECAM1*, *VWF*) 64 lymphocyte (*CCL5*, *CD7*), 405 macrophage (*SDS*, *FCER1G*), 48 mast cell (*TPSB2*, *TPSAB1*) and 36 schwann cell (*SOX10*, *CDH19*) DEGs (FDR<0.1) **(Fig. 1f and Extended Data Fig. 1g)**. Using cell type-specific DEGs, we next identified upregulated cell type-specific pathways across sex and race using the gene ontology database (FDR<0.2).^29^ Endocrine cells were enriched in peptide hormone secretion independent of sex and race **(Fig. 1g and 1h)**. Other cell types showing upregulated cell-type specific pathways included cycling endocrine cells (mitotic cell cycle transition, organelle fission), ductal cells (organic anion transport, branching morphogenesis), acinar cells (digestion, alcohol metabolism), quiescent stellate cells (collagen fibril organization, muscle cell differentiation), activated stellate cells (cell proliferation, cell chemotaxis), endothelial cells (endothelial cell migration, angiogenesis), lymphocytes (immune receptor signaling, T-cell selection), macrophages (antigen processing and presentation, cell chemotaxis), mast cells (immune response, mast cell activation) and schwann cells (CNS myelination and axon development) **(Fig. 1g)**. Cell network analysis confirmed segregation of endocrine pathways from exocrine and immune cell type pathways **(Extended Data Fig. 1h)**. Taken together our data demonstrate that canonical gene networks are conserved across endocrine and non-endocrine cell types independent of sex and race **(Fig. 1e-h, Extended Data Fig. 1h)**.

### Sex differences in islet cell transcriptomes from non-diabetic donors predominantly affect sex chromosome genes

We performed two sets of analysis comparing changes in gene expression in biological variables of sex and race across groups. To study transcriptional differences across donors, we generated principal component analysis (PCA) plots of islet ‘pseudo-bulk’ transcriptional profiles across all 52 donors. Donors did not cluster based on sex, race, disease status, or origin of donor **(Fig. 2a)**. We next segregated donors by cell type, and the resulting PCA showed clustering of samples based on cell type **(Fig. 2b)**. Both whole islet ‘pseudo-bulk’ and individual cell type ‘pseudo-bulk’ sample profiles showed no clustering based on sex or race. This suggests that human islets likely do not have major differences in cell type transcriptional profiles across either race or sex.

**Fig. 2:**
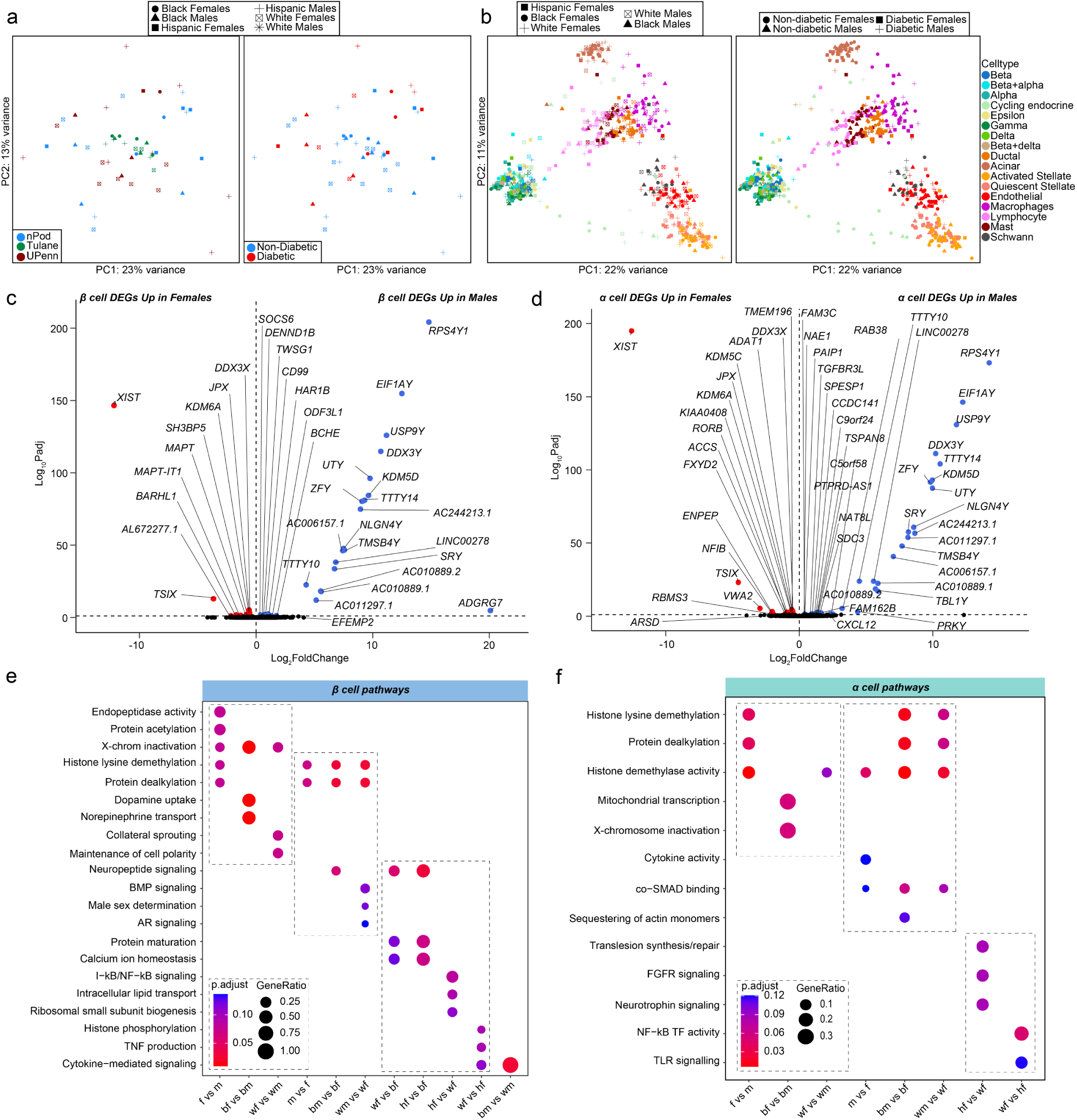
Transcriptional differences across islet β and α cells, highlight enrichment in sex-chromosome genes. **a,** Principal component analysis (PCA) plot of pseudo-bulk transcriptional profiles across all individual donor islets. **b,** PCA plot of pseudo-bulk transcriptional profiles in each cell type across all donors. **c-d,** Volcano plots showing differentially expressed genes (DEGs) across sex in case of non-diabetic: **c,** β cells. **d,** α cells. **e,** GO analysis of all β cell DEGs. **f,** GO analysis of all α cell DEGs. n= 36 non-diabetic and n=16 T2D diabetic donors. DEGs have FDR adjusted q-value <0.1, GO pathways have FDR adjusted q-value <0.2

Focusing on non-diabetic donors, we examined genes with differences in expression between sexes using cell type ‘pseudo-bulk’ analysis. Most sex-associated genes were related to sex chromosomes (FDR<0.1). In β cells, 60% of genes with increased expression in females were linked to the X chromosome and 70% of genes increased in males were linked to the Y chromosome **(Fig. 2c and Extended Data Fig. 2a)**. Similarly, in α cells 50% of male- and 57% of female-enriched genes were linked to the X or Y chromosome, respectively **(Fig. 2d and Extended Data Fig. 2a)**. In α/β cells, X-inactive specific transcript (*XIST*) and lysine demethylase 6A (*KDM6A*) were upregulated in females, while ribosomal protein S4 Y-linked 1 (*RPS4Y1*) and lysine demethylase 5C (*KDM5D*) was upregulated in males **(Fig. 2c and 2d)**. We only observed significant race differences in DEGs between hispanic and white β and α cells **(Extended Data Fig. 2c)**.

Next, we identified sex-specific changes in pathways related to sex chromosome genes using gene set enrichment analyses **(Fig. 2e and Extended Data Fig. 2b)**. Female β cells were enriched for pathways for X-chromosome inactivation and histone lysine demethylation, whereas male β cells were enriched for pathways for Y-chromosome genes, histone lysine demethylation, and male sex determination **(Fig. 2e)**. Female α cells were enriched for histone lysine demethylation, X-chromosome inactivation, and mitochondrial transcription, while male α cells were enriched for histone demethylase activity **(Fig. 2f)**. Similar effects were observed in other cell types **(Extended Data Fig. 2b)**. Race differences in islet cells are shown in **Fig. 2e and 2f** as well as **Extended Data Fig. 2c and 2d**. Of note, black male β cells showed higher cytokine signaling compared to white males, suggesting black male β cells may exhibit a higher inflammatory response **(Fig. 2e).**

### Accessible chromatin landscape across islet cells

To examine the effect of sex on the epigenome, we performed snATAC-seq on all non-diabetic donors of the TUID. To confirm library quality, we filtered and evaluated single nuclei across all 15 donors for TSS enrichment, fragment of reads in promoters, and fragment reads in accessible peaks **(Extended Data Fig. 3a and 3b)**, as well as sample specific sequencing metrics **(Extended Data Fig. 3c and 3d)**. We then clustered the 52,613 filtered profiles resulting in 11 distinct cell clusters which, like gene expression data, were evenly distributed across sex, race, and donor **(Fig. 3a-c)**. To determine the identity of each cluster, we used label transfer to annotate each snATAC-seq cell cluster using our integrated scRNAseq islet cell atlas as a reference. We observed a high degree of correlation between genes with differential accessibility in snATAC-seq and genes with differential expression scRNAseq **(Fig. 3d)**. Cell types also showed a high degree of correlation between RNA expression, chromatin accessibility, and predicted RNA expression **(Extended Data Fig. 3e-g)**. We further examined the cell type annotations using the activity of cell type-specific genes. This validated clusters representing β (*INS-IGF2*), α (*GCG*), δ (*SST*), γ (*PPY*), acinar ductal (*CFTR*), (*PRSS1*), endothelial (*ESM1*), macrophage (*SDS*), stellate *PDGFRA*) and lymphocyte (*CD3D*) cells by comparing gene accessibility with predicted RNA expression **(Fig. 3e and 3f, Extended Data Fig. 3h)**.

**Fig. 3:**
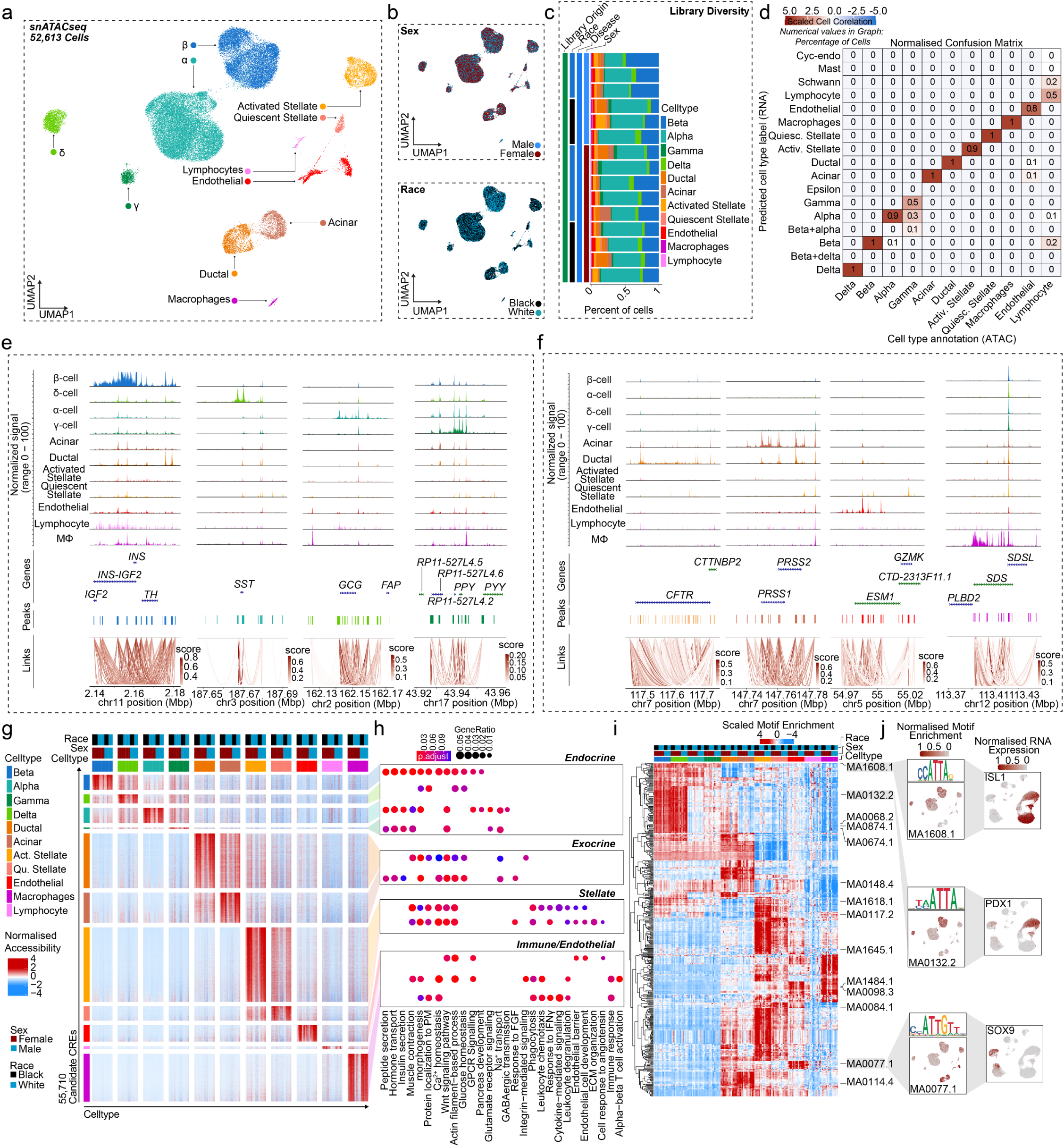
Chromatin accessibility landscape of human pancreatic islet cell types. **a,** UMAP plot denoting integrated clustering of 52,613 single pancreatic islet cells across 11 clustered cell types based on their accessible chromatin profiles, spanning n=15 datasets. Each cluster cell type is denoted by a label and color. **b,** Cell diversified based on sex and race. **c,** Cell distribution stemming from each of the n=15 donors, grouped based on race and sex. **d,** Normalized confusion matrix, showing correlation across cell types based on their cell annotation based on their accessible chromatic profile (x-axis) and predicted cell type label gene expression profile (y-axis). **e,** Aggregated read density profile within a 50-kb window flanking a TSS for selected endocrine marker genes. **f,** Promoter accessibility as in (e) for selected acinar, ductal, endothelial and macrophage genes. **g,** Row normalized chromatin accessibility peak counts for 55,710 candidate cis regulatory elements (CREs) across all 11 cell types. Cells are clustered based on cell type, sex and race. **h,** Gene ontology profiles of differentially active genes based on CREs in **g**. **i,** Row-normalized motif enrichment (ChromVAR) z-scores for the 500 most variable transcription factor motifs, across cell type, sex, and race. Select motifs and corresponding transcription factors are highlighted. **j,** Enrichment z-scores projected onto UMAP coordinates of accessibility for select motifs from **i** (left panel). Normalized RNA expression projected onto UMAP profiles of scRNAseq profiles of islet cells as shown in (Fig. 1a) (right panel). n= 11 non-diabetic donors. Differentially accessible chromatin peak counts have FDR adjusted q-value<0.1, GO pathways have FDR adjusted q-value<0.2

To characterize regulatory programs across each cluster, we identified candidate *cis*-regulatory elements (cCREs) in each cell type resulting in 404,697 total cCREs across all 11 cell types. We next identified cCREs with activity specific to each cell type, resulting in 55,710 cell type-specific cCREs **(Fig. 3g)**. We identified genes in proximity to cell type-specific cCREs, resulting in a list of putative gene targets of cell type-specific regulatory programs. Evaluating these gene sets for enrichment of gene ontology terms revealed cell type-specific processes, and which were similar to those identified in cell type-specific gene expression **(Fig. 3h)**. Using chromVAR^30^, we identified transcription factor (TF) motifs enriched in the accessible chromatin profiles of each cell type using the JASPAR 2020 database.^31^ In-depth analysis of these motifs revealed cell type-specific TF motif enrichment patterns **(Fig. 3i).** For example, we observed enriched motifs for *ISL1* in endocrine cells, *PDX1* in β and δ cells, and *SOX9* in ductal and acinar cells **(Fig. 3i and j)**. These accessible motifs also paralleled cell type specific TF expression in scRNA-seq **(Fig. 3j)**. Similar to previous studies^32–35^, hierarchical motif clustering highlighted that the regulatory programs of β and δ cells are more related, as with α and γ cells **(Fig. 3g)**. Select motifs highly enriched for a cell type (fold enrichment>1.5, −log10 FDR>50) included *PAX4*, *RFX2*, *NKX6-2* and *PDX1* in β cells, *NKX6-2, NKX6-1*, *PDX1*, and *MEOX1* in δ cells, *MAFB*, *FOXD2* and *GATA2-5* in α cells, and *KLF15* and *NRF1* in γ cells **(Extended Data Fig. 3i)**. Non-endocrine cells motif enrichments are also provided in **Extended Data Fig. 3i**.

### Sex differences in chromatin accessibility of islet cells from non-diabetic donors predominantly affects sex chromosomes

To assess sex differences in chromatin accessibility, we identified sex-associated cCREs using logistic regression. As expected, β cells exhibited sex differences in chromatin accessibility at sex chromosome genes including *KDM6A*, *XIST* and *KDM5D* **(Fig. 4a)**. Males exhibited more differentially accessible regions (250 in β, 565 in α) than females (203 in β, 553 in α). Next, we identified genes in a 100 kb proximity to sex-associated cCREs and interrogated their RNA expression. We found that Y-linked genes (*SRY*, *RPS4Y1*, *UTY*, *TTTT14*) in males and X-linked genes (*KDM6A*, *XIST*, *DHRSX*) in females were proximal to sex-associated cCREs **(Fig. 4b)**. Accordingly, when comparing gene expression and cCREs with sex-specific association, we predominantly observed sex-chromosome genes **(Fig. 4c)**. Gene ontology analysis of this subset of genes revealed enrichment in pathways regulating epigenetic control and X chromosome dosage compensation in females, and histone modification in males **(Fig. 4d)**. Notably, the histone demethylase X-linked gene *KDM6A* and the long non-coding RNA *XIST* were more accessible in female islet cells, while the histone demethylase Y-linked gene *KDM5D* was more accessible in males **(Fig. 4e)**. We examined sex differences in TF-specific motif accessibility in α/β cells. Notably, females exhibited a greater number of TF-specific accessible motifs (511 in β, 376 in α) compared to males (33 in β, 74 in α) **(Fig. 4f)**. Upon interrogating differentially expressed TF across cell types, *MAFA*, *SIX3*, *PDX1*, and *RXRG* were upregulated in β cells while *ARX*, *FEV*, *STAT4* and *ISL1* were upregulated in α cells irrespective of sex **(Fig. 4g)**. We applied Pando^36^ to scRNA-seq and snATAC-seq data to infer relationships between target gene expression, TF activation, and TF binding and define gene regulatory networks (GRNs) in male and female β and α cells. The GRNs provide sets of regulated target genes and cCREs for expressed TFs. Irrespective of sex, *MAFA*, *BHLHE41, MEIS2 and MLXIPL* in β cells, and *PAX6* and *SOX5* in α cells, exhibited a high degree of centrality and revealing many associated genes within these TF GRNs **(Fig. 4h)**. In males, *PDX1*, *NKX6-1* and, *RXRG* exhibited higher centrality in β cells, and *ARX* exhibited higher centrality in α cells, compared to females **(Fig. 4h)**.

**Figure 4.**
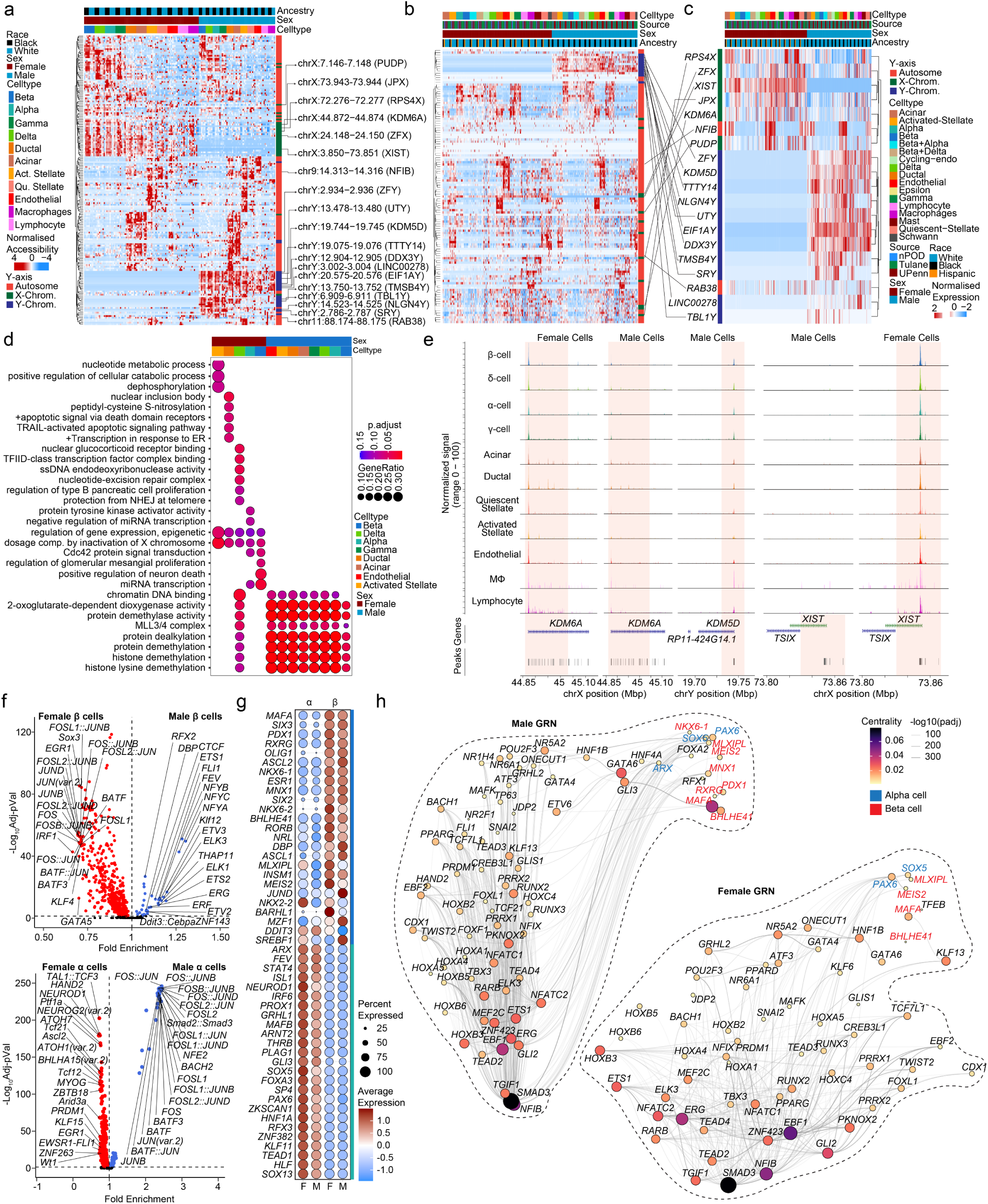
Sex-based enrichment for sex-chromosome gene accessibility in human islet cells. **a,** Row-normalized differentially accessible chromatin peaks across sex and cell-type. *XIST*, *KDM5D* and *KDM6A* are highlighted. **b,** Row normalized expression profiles for genes in a 100kb boundary in proximity to cCREs corresponding to **a** in scRNAseq dataset. **c,** Row normalized expression profiles for the subset of genes corresponding to **b** and differentially expressed genes across sex in scRNAseq dataset. **d,** Gene ontology dot plot showing differential pathways active across multiple cell types based on sex. **e,** Aggregated read density profile within a 50-kb window flanking a TSS for *KDM6A*, *KDM5D* and *XIST.* **f,** Violin plots of differentially accessible motifs identified using ChromVAR in female and male β cells (top) α cells bottom). **g,** Dotplot across sex showing top 25 ranked differentially expressed transcription factors across beta and alpha cells. **h,** Gene regulatory network UMAP embedding of pan-islet transcription factor (TF) activity, based on co-expression, and inferred interaction strength across TFs, for males (left) and females (right). Size/color represent PageRank centrality of each TF. TFs from (g) are highlighted for β (red) and α (blue) cell types. n= 11 non-diabetic donors. Differentially accessible chromatin peak counts have FDR adjusted q-value<0.1, GO pathways have FDR adjusted q-value<0.2.

### Sex and race differences in β cell function

We performed dynamic insulin and glucagon secretion assays in TUID islets for non-diabetic donors. We observed no significant difference in insulin secretion across sex and race using classical insulin secretagogues **(Extended Data Fig. 4a-d)** or an ascending glucose concentration ramp **(Extended Data Fig. 4e-h)**. However, we observed a decreased insulin response to high glucose and IBMX (a phosphodiesterase inhibitor which raises intracellular cAMP) in black male compared to white male islets **(Extended Data Fig. 4a and b)**. We observed no difference of race **(Extended Data Fig. 4i and j)** or sex **(Extended Data Fig. 4k and l)** on α cell function during conditions reflecting hypoglycemia and inhibition of insulin secretion (1.7mM Glucose + 1uM Epinephrine). We also examined the effects of sex and race on islet bioenergetics by quantifying oxygen consumption rate (OCR) **(Extended Data Fig. 4m-p)** and extracellular acidification rate (ECAR) **(Extended Data Fig. 4q-t)** during a glucose challenge in TUID islets. Female islets exhibited greater ATP mediated respiration and coupling efficiency than male islets **(Extended Data Figure 4n and 4p),** suggesting more efficient mitochondria. There was no difference in ECAR between male and female islets.

### Dysregulation of β and α cell transcriptomes from non-diabetic compared with T2D donors suggests sex differences in T2D pathogenesis

We examined the effect of sex on islet hormone secretion using the HPAP islet perifusion database matched for donors we sequenced in this study. Islets from male and female donors with T2D exhibited decreased insulin secretion in response to high glucose, incretin and KCl compared to islets from non-diabetic donors **(Extended Data Fig. 5a and 5b)**, without evidence for sex difference. T2D islets exhibited no difference in α cell function in hypoglycemic conditions compared to non-diabetic donors **(Extended Data Fig. 5c and 5d)**.

We compared the transcriptional profile of male and female HPAP donors with T2D. In contrast with non-diabetic donors, where most sex-associated genes were related to sex chromosomes **(Fig. 2c and 2d)**, islets from T2D donors exhibited multiple sex-specific differences in DEGs from sex chromosomes and autosomes **(Fig. 5a)**. When comparing DEGs in β and α cells from male and female T2D donors, the largest and most significant changes were restricted to sex-linked genes **(Fig. 5b).** We next compared the transcriptional profile of male and female HPAP donors with T2D to that of non-diabetic TUID and HPAP donors **(Extended Data Fig. 1a)**. Notably, in comparison of T2D vs. non-diabetic β cells, females exhibited more DEGs from autosomes (721 upregulated and 1164 downregulated) than males (111 upregulated and 99 downregulated), with only 5.2% of DEGs shared across sex **(Fig. 5c and 5d)**. Similarly, in comparison of T2D vs. non-diabetic α cells, females exhibited more DEGs from autosomes (589 upregulated and 1552 downregulated) than males (14 upregulated and 6 downregulated), with only 0.28% overlap **(Fig. 5c and 5f)**. When comparing T2D vs. non-diabetic donors in other cell types, females also exhibited more autosomal DEGs than males **(Fig. 5c)**. We determined enrichment of gene ontology terms in these genes, and female β and α cells exhibited reduced mitochondrial function and respiration pathways in T2D **(Fig. 5e and 5g)** while male β cells exhibited reduced hormone and insulin secretion pathways in T2D **(Fig. 5e)**. Enrichment of ontology terms for other islet cells in females and males are shown in **Extended Data Fig. 6**.

**Figure 5.**
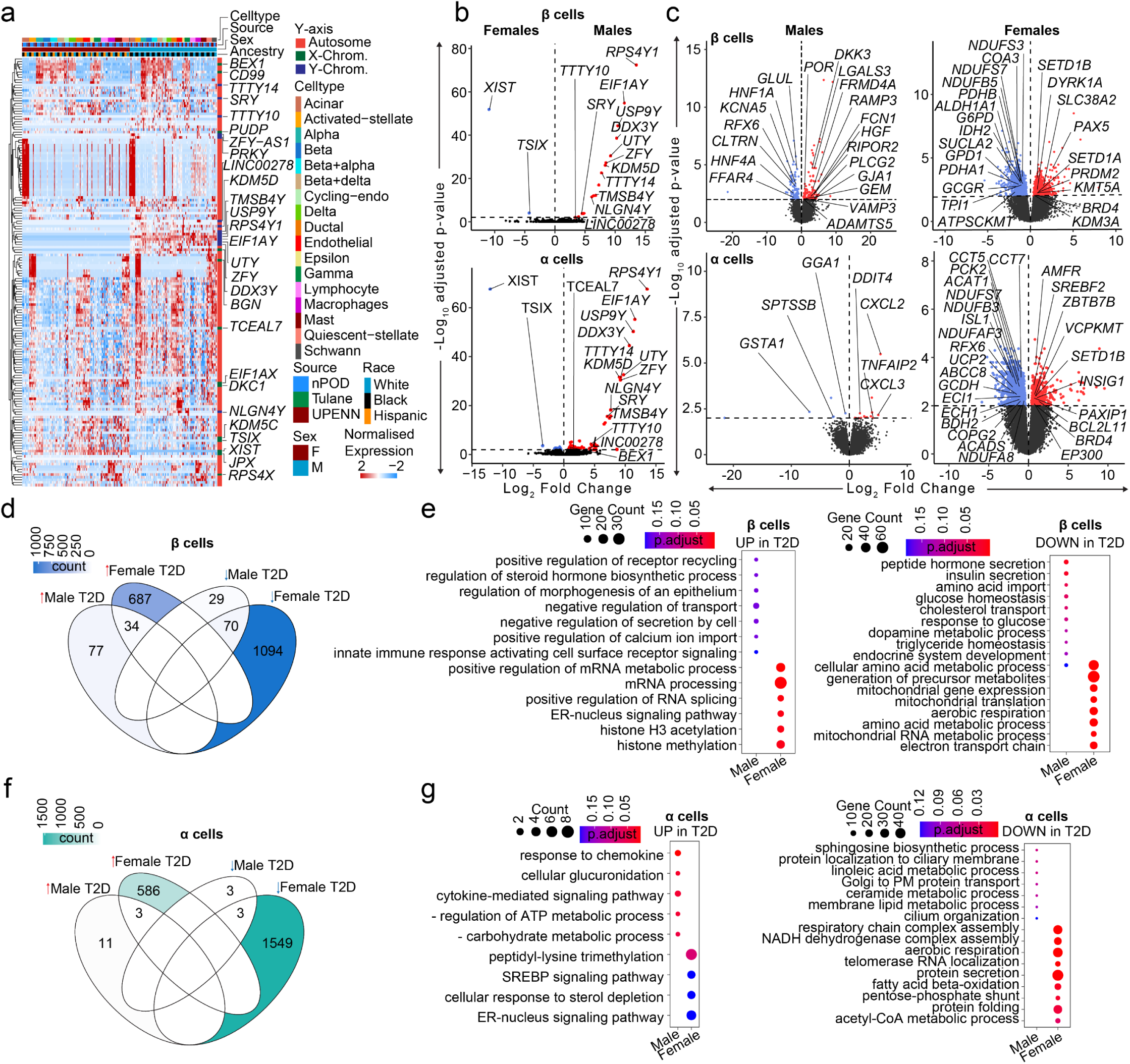
Transcriptional differences in T2D compared to non-diabetic endocrine cells. **a,** Heatmap of DEGs across T2D donors. **b,** Violin plots showing DEGs across male and female T2D β/α cells. **c,** Violin plots showing DEGs across β/α cells when diabetic donors are compared to non-diabetic controls across sex. **d,** Venn diagram showing DEGs across different sex-disease comparisons in case of β cells. Color denotes the number of genes. **e,** Gene ontology dotplot for upregulated and downregulated pathways for β-cell DEGs. **f,** Venn diagram showing DEGs across different sex-disease comparisons in case of α cells. Color denotes the number of genes. **g,** Gene ontology dotplot for upregulated and downregulated pathways for α-cell DEGs. n= 36 non-diabetic and n=16 T2D diabetic donors. DEGs have FDR adjusted q-value<0.01, GO pathways have FDR adjusted q-value<0.2

## Discussion

Our study provides a single cell atlas of sex-specific genomic differences in pancreatic islet cell types in subjects with and without T2D. In non-diabetic islet cells, sex differences in sex-linked genes predominate. In females, *XIST* and its negative regulator *TSIX* are upregulated across all islet cells, suggesting a role of X-chromosome dosage compensation^37^ in human islet function. Similarly, the Y-linked ubiquitin specific peptidase *USP9Y*^38^ and S4 ribosomal protein *RPS4Y1*^39^ genes are expressed exclusively in all male cells, also suggesting a role for these genes in male islet function. Most genes on one X chromosome of XX cells are silenced in development through X chromosome inactivation by *XIST*, thus normalizing X chromosome genes dosage between sexes. However, some X chromosome genes escape inactivation and are expressed from both alleles in XX cells.^40,41^ These “X-escape genes” are conserved between mouse and humans, and several are epigenetic remodelers that promote histone modification to regulate genome access to transcription factors. For example, the histone demethylase *KDM6A* escapes X inactivation^42^ and was more accessible and expressed in female β and α cells. *KDM6A* promotes sex differences in T cell biology.^43^ Similarly, *KDM5D* is only expressed from the male Y chromosome and was overexpressed in male β and α cells. *KDM5D* drives sex differences in male osteogenesis, cardiomyocyte, and cancer.^44–47^ Thus, sex differences in expression of chromatin remodelers like KDM6A or KDM5D may influence sex-specific chromatin access to transcription factors promoting sex differences in islet function. Consistent with this possibility, we observed a five-to-ten-fold greater number of transcription factor-specific accessible motifs in female compared to male α and β cells.

Non-diabetic female islets exhibited greater ATP-mediated respiration and coupling efficiency than those of males, which is consistent with females’ mitochondria having greater functional capacity.^48,49^ In contrast, female β cells from T2D donors showed reduced activation of pathways enriched in mitochondrial function compared to female β cells from non-diabetic donors, which was not observed in male β cells. This suggests that in the transition from normoglycaemia to T2D, female β cell develop greater mitochondrial dysfunction than those of males.^50^ Sex hormones may explain these differences, as estrogen and androgen receptors affect mitochondrial function in female and male β cells.^51,52^ However, since differences between islets from non-diabetic and T2D donors were present outside of the *in vivo* hormonal environment, cell autonomous factors, such as the sexually dimorphic sex chromosomes genes described above are more likely to be involved in these differences.

We find little evidence of differences across race, although inflammatory cytokine signaling was increased in black male β cells via *IL18*, a cytokine implicated in diabetes, obesity, and metabolic syndrome.^53–55^ In addition, non-diabetic black male islets exhibit decreased cAMP-stimulated insulin secretion compared to white male islets. This is reminiscent of ketosis-prone diabetes, a form of T2D mostly observed in males of sub-Saharan African descent with severe β cell failure.^3–5^

A key aspect of our study is the use of ‘pseudo-bulk’ profiles aggregated per cell type in each sample. Collapsing cell profiles by sample enables to effectively control for pseudo-replication due to cells being sampled from a fixed number of donors, whereas treating each cell from the same cluster as an independent observation leads to inflated p-value and spurious results. This approach has demonstrated high concordance with bulk RNA-seq, proteomics and functional gene ontology data.^56,57^ We applied a hypergeometric statistical model using ‘pseudo-bulk’ count data correcting for library composition bias and batch effects in the scRNA-seq.^26^ This approach has enabled us to recapitulate biological ground truth, where we demonstrate high concordance between accessible chromatin and associated active genes across human islet cells.

In conclusion, this study establishes an integrated accessible chromatin and transcriptional map of human islet cell types across sex and race at single cell resolution, reveals that sex-specific genomic differences in non-diabetic individuals predominantly through sex chromosome genes, and reveals genomic differences in islet cell types in T2D which highlights mitochondrial failure in females.

### Limitations of the study

Despite the inclusion of seven black donors (Tulane dataset) to promote genetic diversity, our study is limited by the small number of donors. Future extramural funding for the inclusion and study of diverse genetic datasets is essential. Another key consideration is library composition bias owing to targeted islet sequencing, which is not a representation of all pancreatic cells, cell subtypes, or spatiotemporal domains.^58,59^ Even after utilizing a stringent ambient RNA correction methodology, invariably residual contaminant RNA can be observed across cells. Emphasis is given on generating tools to adjust for ambient RNA particularly in case of pancreatic cells containing high expression of genes such as INS and PRSS1.

## Acknowledgments

This work was supported by National Institutes of Health grants DK074970 (F.M.-J.), DK105554 (K.J.G), HG012059 (K.J.G), DK114650 (K.J.G), DK120429 (K.J.G), U.S. Department of Veterans Affairs Merit Award BX005812 (F.M.-J.), and the Tulane Center of Excellence in Sex-Based Biology & Medicine (F.M.-J.). The preparation of human pancreatic islets provided by the Integrated Islet Distribution Program (IIDP) (RRID:SCR_014387) at City of Hope were funded by NIH grant 2UC4DK098085.

## Author Contributions

MMFQ, KS, SSVPS, SH, designed and/or performed/analyzed experiments, MMFQ, RME, PK designed and/or performed/analyzed computational experiments, MMFQ and RME prepared the final figures and wrote/edited the manuscript. PVK, SED, JK, KJG, provided reagents and analyzed experiments. F.M.-J. designed the study, analyzed the data, and wrote and revised the manuscript. All authors reviewed and edited the manuscript and accepted the final version.

## Declaration of interests

The authors declare no conflict of interest

## Lead contact

Further information and requests for resources and reagents should be directed to and will be fulfilled by the lead contact, Franck Mauvais-Jarvis.

## Materials availability

This study did not generate any new materials.

## Data and code availability

- Single cell RNA and single nuclei ATAC sequencing data has been deposited at GEO (deposition will be made public upon publication), All data reported in this paper will be shared by the lead contact upon request.
- A description of coding environments required to reproduce scRNAseq analysis in this paper are outlined in: https://github.com/FMJLabTulane/sex_regulome_pancreas
- Any additional information required to reanalyze the data reported in this paper is available from the lead contact upon request.

## Human pancreatic islets

De-identified human pancreatic islets from fifteen male and female donors were obtained from PRODO Laboratories Inc, and the Integrated Islet Distribution Program (IIDP). Islets were left in culture at 37°C in a humidified incubator containing 5% CO_2_ overnight before any experiments were performed. Islets were cultured in phenol-red free RPMI medium (Gibco) containing 11mM glucose, supplemented with 10% Charcoal Stripped FBS (Invitrogen), HEPES (10mM; Gibco), Sodium Pyruvate (1mM; Gibco), β-mercaptoethanol (50µM; Invitrogen), GlutaMAX (2mM; Gibco) and Penicillin-Streptomycin (1x; Gibco).

## Studies involving Human cadaveric tissue

Samples originate from de-identified cadaveric donors and are institutional review board exempt.

## Measurement of insulin secretion in perifusion

Perifusion experiments were performed in Krebs buffer containing 125mM NaCl, 5.9mM KCl, 1.28mM CaCl2, 1.2mM MgCl2, 25mM HEPES, and 0.1% bovine serum albumin at 37°C using a PERI4-02 machine (Biorep Technologies). Fifty hand-picked human islets were loaded in Perspex microcolumns between two layers of acrylamide-based microbead slurry (Bio-Gel P-4, Bio-Rad Laboratories). For experiment 1, cells were challenged with either low or high glucose (5.6mM or 16.7mM), IBMX (100μM), epinephrine (1μM) or potassium chloride (20mM) at a rate of 100µL/min. After 60 minutes of stabilization in 5.6mM glucose, cells were stimulated with the following sequence: 10min at 5.6mM glucose, 30min at 16.7mM glucose, 15min at 5.6mM glucose, 5min at 100μM IBMX + 16.7mM glucose, 15min at 5.6mM glucose, 5min at 1μM epinephrine + 1.7mM glucose, 15min at 5.6mM glucose, 15min at 20mM KCl + 5.6mM glucose, and 15min at 5.6mM glucose. In case of experiment 2, islets were challenged with either low or graded high concentrations of glucose (2, 5, 11 or 20mM) or potassium chloride (20mM) at a rate of 100μL/min. After 60min of stabilization in 2mM glucose, islets were stimulated in the following sequence: 10min at 2mM glucose, 10min at 7mM glucose, 10min at 11mM glucose, 10min at 20mM glucose, 15min at 2mM glucose, 10min at 20mM KCl + 2mM glucose, 10min at 20mM KCl + 11mM glucose and, 10min at 2mM glucose. Samples were collected every minute on a plate kept at <4°C, while the perifusion solutions and islets were maintained at 37°C in a built-in temperature controlled chamber. Insulin and glucagon concentrations were determined using commercially available ELISA kits (Mercodia). Total insulin and glucagon release was normalized per total insulin or glucagon content respectively using a human insulin or glucagon ELISA kit (Mercodia).

For samples used as a part of the HPAP dataset, sample metadata and perifusion data were downloaded from the HPAP website: https://hpap.pmacs.upenn.edu/, for samples used as a part of this study. Data were organized based on insulin and glucagon secretion where available and plotted across sex.

## Bioenergetics

Islets were washed once with assay buffer (made from Agilent Seahorse XF Base Medium supplemented with 3mM glucose and 1% charcoal striped FBS). Around 150 islets were transferred to each well of Seahorse XF24 Islet Capture Microplate (Agilent) and were incubated in assay buffer at 37 °C for 60 minutes before being transferred to Agilent Seahorse XFe24 Analyzer. Islets were maintained in the assay medium throughout the experiment, while oxygen consumption rate (OCR) and extracellular acidification rate (ECAR) were measured at basal (3 mM), glucose-stimulated level (20 mM) and after addition of oligomycin, carbonyl cyanide-4 (trifluoromethoxy) phenylhydrazone (FCCP), rotenone/antimycin according to manufacturer’s instructions.

## Single cell RNA indexing and sequencing

Human islets (500 IEQ per condition) were cultured overnight in a humidified incubator containing 5% CO2 at 37°C. Islet cells were then dispersed using TrypLE (Thermofischer), and immediately evaluated for viability (90.61±3.04%) by Cellometer Automated Cell Counter (Nexcelom Bioscience) prior to single cell RNAseq library preparation. For 10x single cell RNAseq library preparation, 5000-6500 individual live cells per sample were targeted by using 10x Single Cell 3’ RNAseq technology provided by 10x Genomics (10X Genomics Inc). Briefly, viable single cell suspensions were partitioned into nanoliter-scale Gel Beads-In-EMulsion (GEMs). Full-length barcoded cDNAs were then generated and amplified by PCR to obtain sufficient mass for library construction. Following enzymatic fragmentation, end-repair, A-tailing, and adaptor ligation, single cell 3’ libraries comprising standard Illumina P5 and P7 paired-end constructs were generated. Library quality controls were performed by using Agilent High Sensitive DNA kit with Agilent 2100 Bioanalyzer (Agilent) and quantified by Qubit 2.0 fluorometer (ThermoFisher). Pooled libraries at a final concentration of 750pM were sequenced with paired end single index configuration by Illumina NextSeq 2000 (Illumina).

## Single cell gene expression mapping

For the Tulane dataset we utilized CellRanger v4.0.0 software using the [-mkfastq] command to de-multiplex FASTQ data. Reads were mapped and aligned to the human genome (10X genomics pre-built GRCh38-2020-A Homo sapiens reference transcriptome assembly) with STAR (95.33±0.75% of reads confidently mapped to the human genome).^60^ Subsequently, final digital gene expression matrices and c-loupe files were generated for downstream multimodal analysis. In case of the HPAP dataset we isolated data processed as described previously (nPod data: 87.91±11.56 and UPenn 90.62±5.44% of reads map confidently to genome).^26^ Cellranger identified 75,619 (Tulane), 73,472 (nPOD) and 52,357 (UPenn) correctly allocated barcodes (cells), having 78,584±40,590 (Tulane), 130,993±289,368 (nPOD), 63,949±29,598 (UPenn) reads/cell and 26,866±680 (Tulane), 24,739±8983 (nPOD), 24,183±1254 (UPenn) genes/cell.

## Preliminary filtering and S4 R object creation

We deployed Seurat v4.3.0^61,62^ scripts to perform merging, thresholding, normalization, principal component analysis (linear dimensionality reduction), clustering analysis (non-linear multidimensional reduction), visualization and differential gene expression analysis. Cells having total mitochondrial RNA contribution beyond 20% were eliminated from the analysis, along with cells expressing less than 500 or greater than 8000 total genes.

## Ambient RNA correction and doublet annotation

In droplet based scRNAseq technologies, extracellular RNA from cells with compromised membrane integrity contaminates single cell libraries.^56^ This remains a challenge for pancreatic cells, as endocrine and exocrine cells are rich in select secreted RNA species. We used SoupX 1.6.1^63^ on raw feature barcode matrices correcting for ambient RNA across all 52 donors. Raw counts were corrected using SoupX and rounded to the nearest integer. As the TUID is not doublet corrected, we utilized DoubletFinder v2^64^ expecting 5% doublets, eliminating them from the dataset.

## Data normalization and clustering

SoupX corrected matrices were metadata annotated, and geometrically normalized (log10) at a scale factor of 10,000. The variance stabilization method (vst) method was used to find 2000 most variable features, which were later used for scaling and principal component analysis (PCA) using 20 components. and dimensions (UMAP). We batch corrected the datasets using Harmony 0.1.1^65^, using donor library identity, 10X genomics chemistry (v2 or v3) and tissue source (Tulane, nPOD or UPenn) as covariates in the batch model. Uniform manifold approximation and projection (UMAP) and neighbors were calculated using Seurat v4.3.0.^61,62^ Finally we hyperclustered data using a Leiden algorithm at a resolution of 6. We observed poor quality cells to remain in the dataset (low relative total RNA and gene counts yet within threshold), and excluded these from the analysis, and performed re-clustering as described above. Finally, we assigned identities to clusters based on pancreatic cell specific gene sets^28,58^, resulting in 17 discrete clusters, totaling 141,739 high quality cells.

## Cell type specific marker genes

Statistical approaches to define DEGs across cell types using aggregated “pseudobulked” RNA count data, out-perform single cell DEG models^56,57,66^. Infact, pseduobulk DEG methods demonstrate the highest Mathews Correlation Coefficient, a balanced machine learning performance testing model, capable of evaluating models classifying binary data.^66,67^ Therefore, we performed an unbiased differential analysis of cell cluster-specific marker genes using the [FindAllMarkers] function in Seurat. We employed DESeq2 v1.36.0^68^ to perform DEG testing, where a cluster must express a gene in at least 25% of cells, have a 2x fold difference, and a Benjamini-Hochberg FDR adjusted p-value < 0.01 (α = 1%). Aggregated counts were compared across cell types and donors.

## Sex, race, and disease type specific marker genes

Based on facts outlined above, we employ a previously described statistical model^26^ using DESeq2 v1.36.0^68^ to evaluate statistical differences across human islet cell types based on race, sex and disease, metadata profiles across donors. A DEG is defined as a gene having a Benjamini-Hochberg adjusted p-value < 0.1 (α = 10%).

## Single nuclear assay for transposase-accessible chromatin indexing and sequencing

Human islets (500 IEQ per condition) were cultured overnight in a humidified incubator containing 5% CO2 at 37°C. Islet cells were then dispersed using TrypLE (Thermofischer), and immediately evaluated for viability (90.61±3.04%) by Cellometer Automated Cell Counter (Nexcelom Bioscience) prior to single nuclei ATAC library preparation. Nuclei were isolated based on the 10X genomics Nuclei isolation protocol (CG00169 Rev D) with some modifications. We observe that the usage of 0.5ml tubes yields superior nuclei collection. Furthermore, we optimize based on a sample-to-sample basis the time for cell lysis (3-5min). The final lysis buffer concentration for Nonidet P40 was 0.15% over the 0.1% recommendation. Finally, in addition to the final wash with wash buffer, we perform a final wash with the 10X Genomics Nuclei Buffer (PN-2000153/2000207). Nuclei are always kept < 0°C, visually inspected for integrity and quality using a viability dye, prior to library prep which was performed within 30min. Briefly, 5,000-6,500 isolated nuclei were incubated with a transposition mix to preferentially fragment and tag the DNA in open regions of the chromatin. The transposed nuclei were then partitioned into nanoliter-scale Gel Bead-In-emulsions (GEMs) with barcoded gel beads, a master mix, and partition oil on a chromium chip H. Upon GEM formation and PCR, 10x barcoded DNA fragments were generated with an Illumina P5 sequence, a 16nt 10x barcode, and a read 1 sequence. Following library construction, sequencing-ready libraries were generated with addition of P7, a sample index, and a read 2 sequence. Quality controls of these resulting single cell ATAC libraries were performed by using Agilent High Sensitive DNA kit with Agilent 2100 Bioanalyzer (Agilent) and quantified by Qubit 2.0 fluorometer (ThermoFisher). Pooled libraries at a final concentration of 750pM were sequenced with paired-end dual indexing configuration by Illumina NextSeq 2000 (Illumina) to achieve 40,000-30,000 read pairs per nucleus.

## Single nuclei accessible chromatin mapping

We utilized CellRanger ATAC v1.2.0 software using the [-mkfastq] command to de-multiplex FASTQ data. Reads were mapped and aligned to the human genome (10X genomics pre-built GRCh38-2020-A Homo sapiens reference transcriptome assembly) with STAR (70.70±11.46% of reads confidently mapped to the human genome).^60^ Cellranger identified 84,741 correctly annotated barcodes (cells), having an average transcriptional start site (TSS) enrichment score of 6.27±1.38 and 73.55±6.78% fragments overlapping peaks/sample. We then utilized Signac’s peak calling tool to call peaks on our dataset using MACS2.^69^ We utilize the [CallPeaks()] function to annotate accessible peaks using MACS2.

## Preliminary filtering and S4 R object creation

We deployed Seurat v4.3.0^61,62^ coupled with Signac v1.10.0^70^ scripts to perform merging, thresholding, normalization, principal component analysis (linear dimensionality reduction), clustering analysis (non-linear multidimensional reduction), visualization and differential gene expression analysis. Cells having a TSS enrichment score of < 2, peak region fragments less than 2000 or more than 20,000 counts, percentage reads in peaks < 30%, blacklist ratio > 0.05, nucleosome ratio > 4 and, fraction reads in promoters < 0.2 were eliminated from the analysis.

## Doublet annotation

It is increasingly challenging to detect multiplets in droplet based snATAC data, owing to sparsity and low dynamic range. We employed AMULET^71^ within the scDblFinder v1.10.0^72^ R package on raw fragment barcode matrices correcting for all 15 donors, using the authors recommendations.

## Data normalization and clustering

We used a unified set of peaks across all 15 datasets, annotating genes using EnsDb.Hsapiens.v86.^73^ We estimated gene activity using Signac’s GeneActivity function, by extracting gene coordinates and extend them to include the 2 kb upstream region, followed by geometric normalization (log10). We next performed non-linear multidimensional reduction using term frequency-inverse document frequency (TF-IDF) weighted peak counts transformed to binary data. Weighted data was reduced to 30 dimensions using RunSVD function. We batch corrected the datasets using Harmony 0.1.1^65^ using 30 nearest neighbours, using donor library identity as a covariate in the batch model. The first singular value decomposition (SVD) component correlated with read depth and was eliminated from UMAP projection dimensionality reduction, and SLM^74^ clustering, based on recommendations provided in Signac.

Upon performing iterative clustering and after removing low quality cells, we end up with 52,613 nuclei having 255,194 peak features spanning 11 clusters. We classified clusters based on described gene activities across islet cells,^32^ followed by validating identity with label transfer, from our RNAseq atlas dataset using the FindTransferAnchors function. Finally, we stored an additional modeled predicted RNA expression matrix within the snATAC object using the TransferData function.

## Cell type specific marker genes

To evaluate differentially accessible regions (DARs) we used a Wilcoxon rank sum test comparing a cluster of cells against all other clusters, defining DARs as those peaks expressed in atleast 5% of cells, having a foldchange > 2, Benjamini-Hochberg FDR adjusted pvalue < 0.05 (α = 5%) and restricting to those peaks that are within a 100kb window of a gene.

## Sex, race, and disease type specific marker genes

In order to evaluate population wide differences, we employed the similar model utilized for scRNAseq.^26^ A DAR is defined as a peak having a Benjamini-Hochberg adjusted p-value < 0.1 (α = 10%).

## Single-Cell motif enrichment

We used chromVAR v1.22.1^30^ to estimate transcription factor motif enrichment z-scores across all cells. We used a peak by cell sparse binary matrix correcting for GC content bias based on the hg38 genome (BSgenome.Hsapiens.UCSC.hg38). We use the non-redundant JASPAR 2020 core vertebrate motif database^75^ calculating bias-corrected deviation z-scores across single cells. We then calculated average transcription factor motif enrichment z-scores across single cells in a cluster. We used aggregate cell average z-scores to evaluate differentially accessible motifs (DAMs) across clusters, using a Benjamini-Hochberg FDR corrected p-value < 0.05.

## Gene set enrichment and pathway analysis

In order to perform gene set enrichment analysis (GSEA)^76^, we downloaded the entire molecular signatures database (MSigDB) v3^76,77^ for C5 human gene ontological terms, using clusterProfiler v4.4.4^78^ or using an R based deployment (https://github.com/wjawaid/enrichR) of EnrichR.^79^ We subset the C5 database, restricting terms to biological processes and perform functional pathway annotation using the compareCluster function. We define a pathway to be statistically significant at a Benjamini-Hochberg FDR adjusted p-value < 0.2 (α = 20%). We performed functional pathway mapping using the cnetplot function.

## Gene regulatory network analysis

In order to infer gene regulatory networks (GRNs) we utilized Pando^36^ while using the predicted RNA expression profile and MACS2 components of our snATAC dataset while interrogating TFs for which motifs exist. The coefficients of Pando’s model highlight a quantified measure of interaction across cCRE-TF pair and a downstream target gene, resulting in a regulatory graph which can be plotted using non-linear multidimensional reduction.

## Notes

### Competing Interest Statement

The authors have declared no competing interest.

### Summary of Updates

Corrections have been made to Grant funding identification numbers which were incorrect in previous versions.

