## Extended Data for "Single cell regulatory architecture of human pancreatic islets suggests sex differences in β cell function and the pathogenesis of type 2 diabetes"

[1]Section of Endocrinology and Metabolism, John W. Deming Department of Medicine, Tulane University School of Medicine, New Orleans, LA, USA
[2]Southeast Louisiana Veterans Health Care System, New Orleans, LA, USA
[3]Tulane Center of Excellence in Sex-Based Biology & Medicine, New Orleans, LA, USA
[4]Biomedical Sciences Graduate Program, University of California, San Diego, La Jolla, CA, USA
[5]Department of Pediatrics, University of California, San Diego, La Jolla, CA, USA
[6]Center for Translational Research in Infection and Inflammation, John W. Deming Department of Medicine, Tulane University School of Medicine, New Orleans, LA, USA
[7]Department of Pharmacology, Tulane University School of Medicine, New Orleans, LA, USA
[8]Department of Pediatrics, Tulane University, School of Medicine, New Orleans, LA, USA

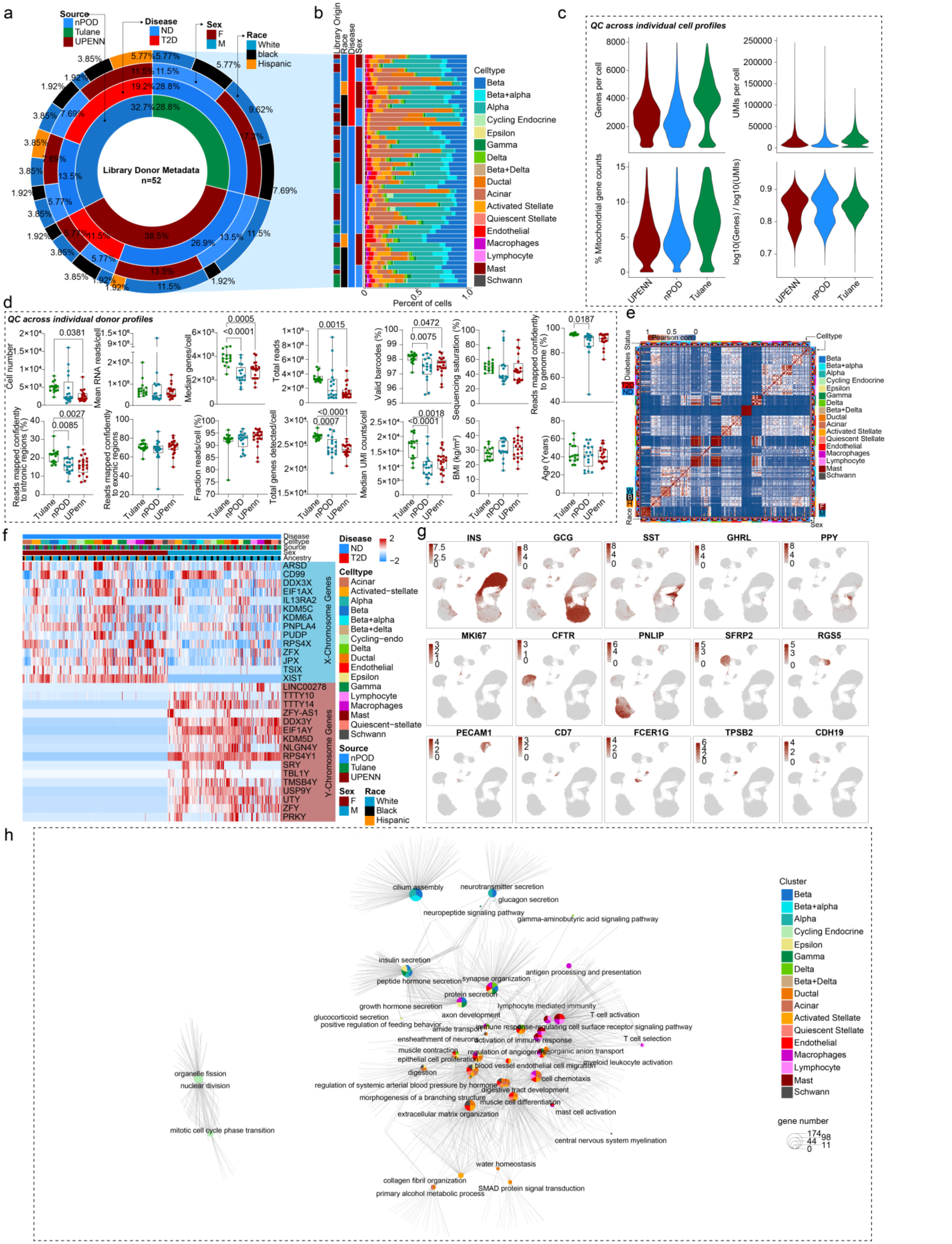

**Extended Data Fig.1 Donor metadata, scRNAseq quality control metrics and network analysis of all non-diabetic cell cells.**

**a,** Sunburst plot showing donor metadata of scRNAseq datasets used in this study. n=52 donors from Tulane University and the Human Pancreas Analysis Program (HPAP). Metadata is organized into source, disease status, sex, and race across all donor profiles. **b,** Percentage distribution across all 17 cell clusters for each of the n=52 donors used in this study. Cells are colored based on a key (legend) and metadata is organized based on information shown in **(a)**. **c,** Violin plots showing genes expressed per cell, unique molecular identifiers (UMIs) per cell, percentage mitochondrial genes and library complexity [log10(number of genes)/log10(number of UMIs)] across each of the donor sources. **d,** Quality control metrics for cell number, mean RNA reads/cell, median genes/cell, total reads, valid barcodes, sequencing saturation, percentage of reads mapped confidently to the reference genome, reads mapped confidently to intronic regions, reads mapped confidently to exonic regions, fraction of reads/cell, total genes detected/cell, median UMI counts/cell, basal metabolic index and age, across all donors split on the basis of source. **e,** Pearson's correlation between normalized read coverage within a merged set of RNAseq expression profiles across 52 donors. Annotation bars highlight the sex, race, cell type and diabetes status of the donors. **f,** Heatmap showing normalized aggregated RNAseq expression profiles for select Y and X linked chromosome genes across race, sex, source, celltype and disease. **g,** UMAP plots showing RNAseq expression profiles for select cognate genes across each cell type specific cell cluster. **h,** Gene ontology network analysis showing regulated pathways across all non-diabetic islet cells. Size of circle equivalent to gene contribution. **(d)** $p<0.05$ is significant **(h)** FDR adjusted $p<0.2$ is significant. n = 52 donors

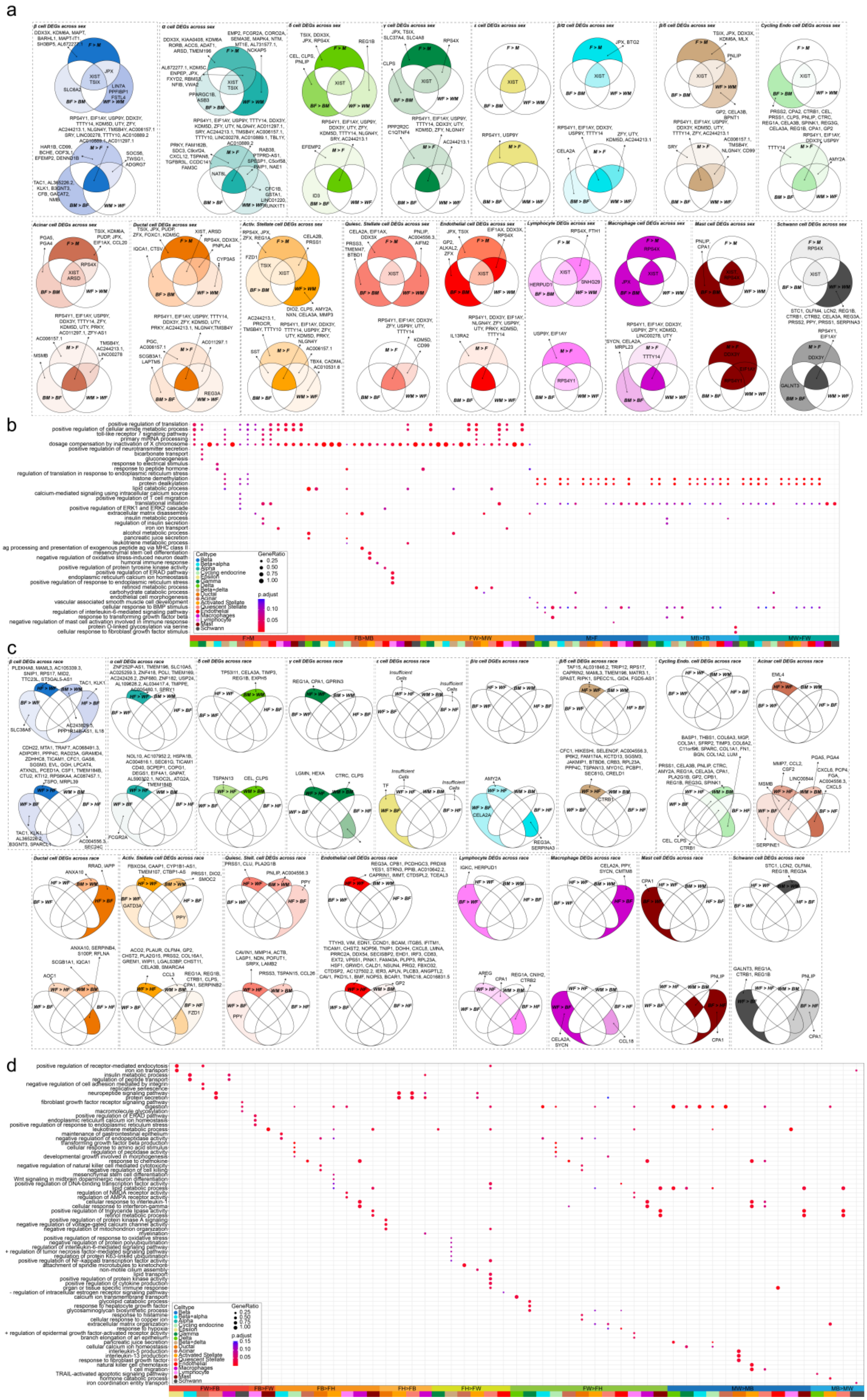

**Extended Data Fig.2 Differentially expressed genes and pathways in human islet cells across sex and race.**

**a,** Venn diagrams showing differentially expressed genes (DEGs) across sex in case of non-diabetic: beta, alpha, delta, gamma, beta+alpha, beta+delta, cycling endo, acinar, ductal, activated stellate, quiescent stellate, endothelial, lymphocyte, macrophage, mast and schwann cells. **b,** Gene ontology (GO) analysis of celltypes other than α/β shown in **(a)**. **c,** Venn diagrams showing differentially expressed genes (DEGs) across ancestry for cells highlighted in **(a)**. **d,** Gene ontology (GO) analysis of all celltypes other than α/β shown in **(c)**. DGE: FDR adjusted pval < 0.1 is significant, GO: FDR adjusted pval < 0.2 is significant, n = 36 (non-diabetic).

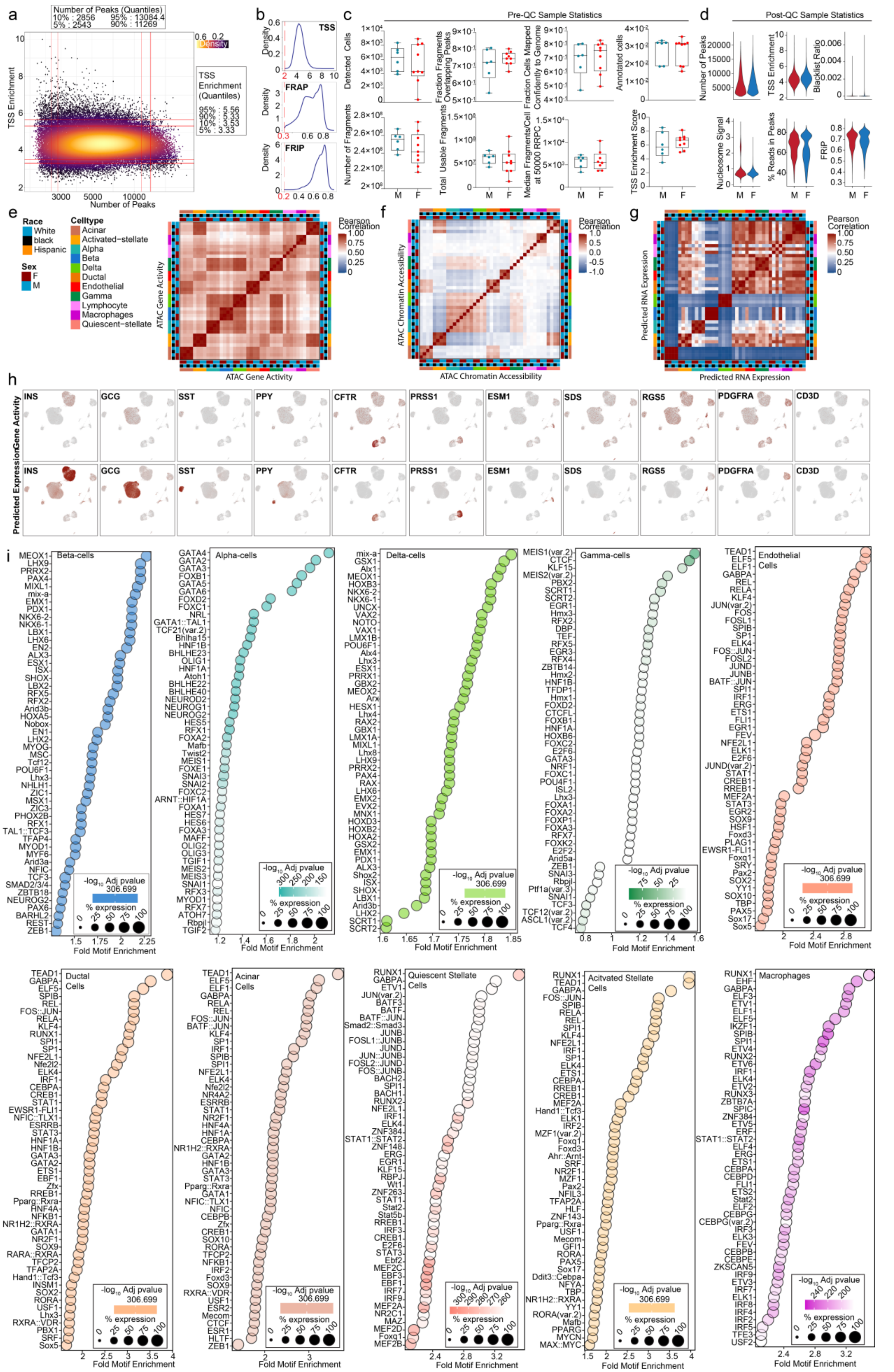

**Extended Data Fig.3 snATACseq quality control metrics and network analysis of all non-diabetic cells.**

**a,** Density scatter showing transcriptional start site (TSS) enrichment on y-axis and number of peak counts on the x-axis. **b,** QC plots showing TSS, fragment reads in accessible peaks (FRAP) and fragment reads in promoters (FRIP) across all donors. Dashed red line signifies cutoff. **c,** QC plots showing metrics for detected cells, fraction fragments overlapping peaks, fraction cells mapped confidently to genome, annotated cells, number of fragments, total usable fragments, median fragments/cell at 50K raw reads/cell (RRPC) and TSS enrichment score across all donors. **d,** Post-filtering QC metrics showing number of peaks, TSS enrichment, blacklist ratio, nucleosome signal, percentage reads in peaks and FRIP across all donors. **e,** Pearson's correlation between aggregated read coverage from snATACseq experiments. **f,** Pearson's' correlation between aggregated gene activity from snATACseq experiments. **g,** Pearson's' correlation between aggregated predicted RNA expression in snATACseq experiments based on mapped expression models from scRNAseq experiments. **h,** Gene activity based on accessibility in 100kb windows flanking select cognate genes across cell clusters (top row). Predicted gene expression based on expression data mapping from scRNAseq atlas (bottom row). **i,** Differential analysis using chromVAR and the JASPAR 2020 motif database showing enriched motifs for beta, delta, alpha, gamma, ductal, acinar, quiescent stellate, activated stellate, endothelial and macrophage cells. DGE: FDR adjusted pval < 0.05 is significantly enriched. n = 15 non-diabetics.

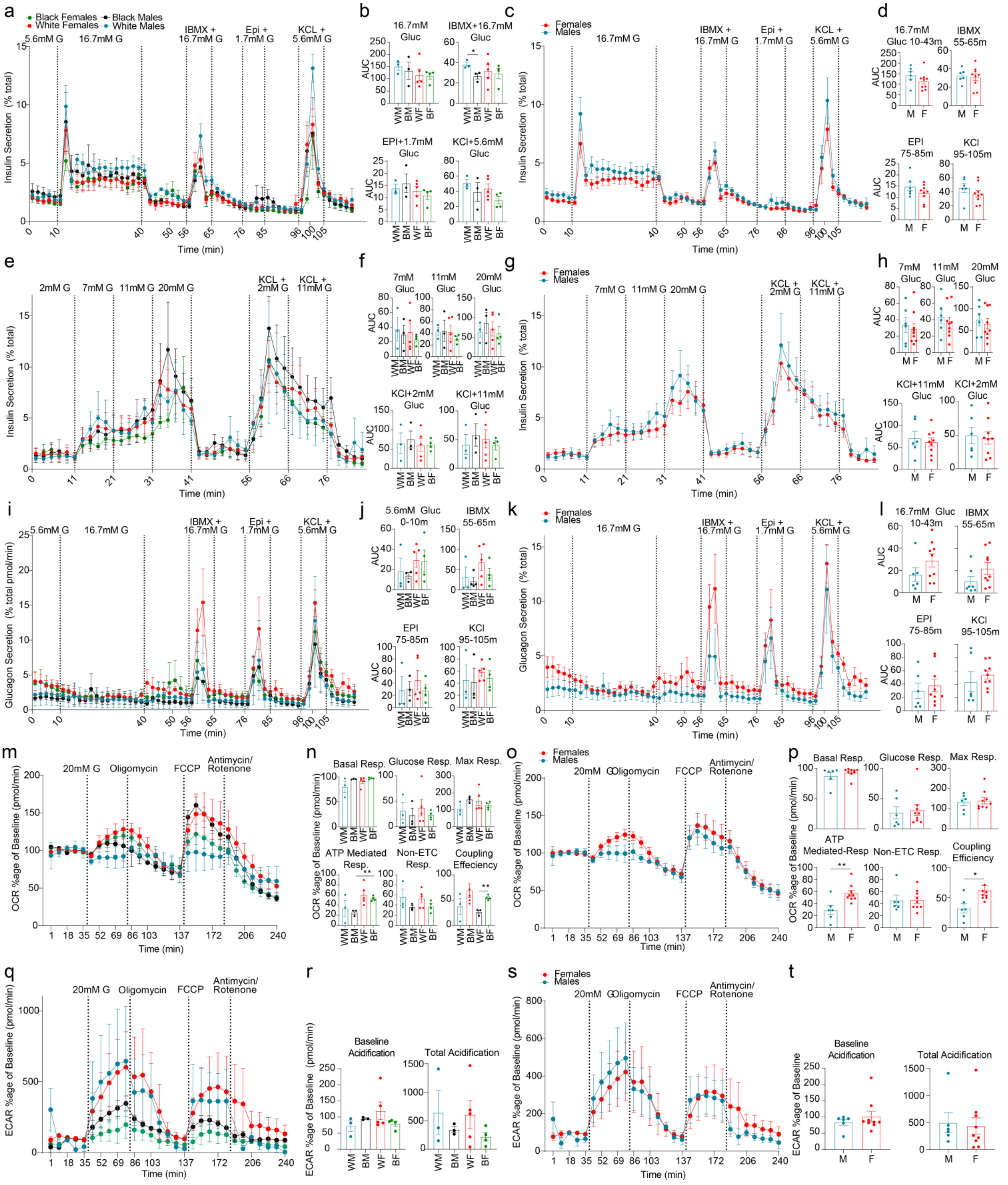

**Extended Data Fig.4 Dynamic insulin and glucagon secretion coupled with islet bioenergetics.**

**a,** Dynamic insulin secretion assay, showing response to 16.7mM glucose, IBMX + 16.7mM Glucose, epinephrine + 1.7mM Glucose and potassium chloride + 5.6mM glucose. Each curve represents secretion normalized to total insulin content across sex and race. **b,** Area under the curve (AUC) measurements for incretin driven insulin secretion measurements outlined in (a). **c,** Dynamic insulin secretion assay, showing response to 16.7mM glucose, IBMX + 16.7mM Glucose, epinephrine + 1.7mM Glucose and potassium chloride + 5.6mM glucose. Each curve represents secretion normalized to total insulin content across sex. **d,** Area under the curve (AUC) measurements for incretin driven insulin secretion measurements outlined in **(b)**. **e,** Dynamic insulin secretion assay using an ascending glucose ramp. Stimulation modes show response to 7mM glucose, 11mM Glucose, 20mM Glucose, potassium chloride + 2mM glucose and, potassium chloride + 11mM glucose. Each curve represents secretion normalized to total insulin content across sex and race. **f,** Area under the curve (AUC) measurements for incretin driven insulin secretion measurements outlined in **(e)**. **g,** Dynamic insulin secretion assay using an ascending glucose ramp. Stimulation modes show response to 7mM glucose, 11mM Glucose, 20mM Glucose, potassium chloride + 2mM glucose and, potassium chloride + 11mM glucose. Each curve represents secretion normalized to total insulin content across sex and race. **h,** Area under the curve (AUC) measurements for incretin driven insulin secretion measurements outlined in **(g)**. **i,** Dynamic glucagon secretion assay, showing response to 16.7mM glucose, IBMX + 16.7mM Glucose, epinephrine + 1.7mM Glucose and potassium chloride + 5.6mM glucose. Each curve represents secretion normalized to total glucagon content across sex and race. **j,** Area under the curve (AUC) measurements for incretin driven insulin secretion measurements outlined in **(i)**. **k,** Dynamic glucagon secretion assay, showing response to 16.7mM glucose, IBMX + 16.7mM Glucose, epinephrine + 1.7mM Glucose and potassium chloride + 5.6mM glucose. Each curve represents secretion normalized to total glucagon content across sex. **l,** Area under the curve (AUC) measurements for incretin driven insulin secretion measurements outlined in **(k)**. **m,** Oxygen consumption ratio for islets across sex and race. **n,** Basal respiration, glucose mediated respiration, maximal (max) respiration, ATP mediated respiration, non-electron transport chain (ETC) respiration and coupling efficiency, across sex and race. **o,** Oxygen consumption ratio for islets across sex. **l,** Basal respiration, glucose mediated respiration, maximal (max) respiration, ATP mediated respiration, non-electron transport chain (ETC) respiration and coupling efficiency, of human islets across sex. **q,** Extracellular acidification rate for islets across sex and race. **r,** Baseline acidification and total acidification of human islets across sex and race. **s,** Extracellular acidification rate for islets across sex. **t,** Baseline acidification and total acidification of human islets across sex. *pval < 0.05, **pval < 0.01 is significant. n = 15 (non-diabetic).

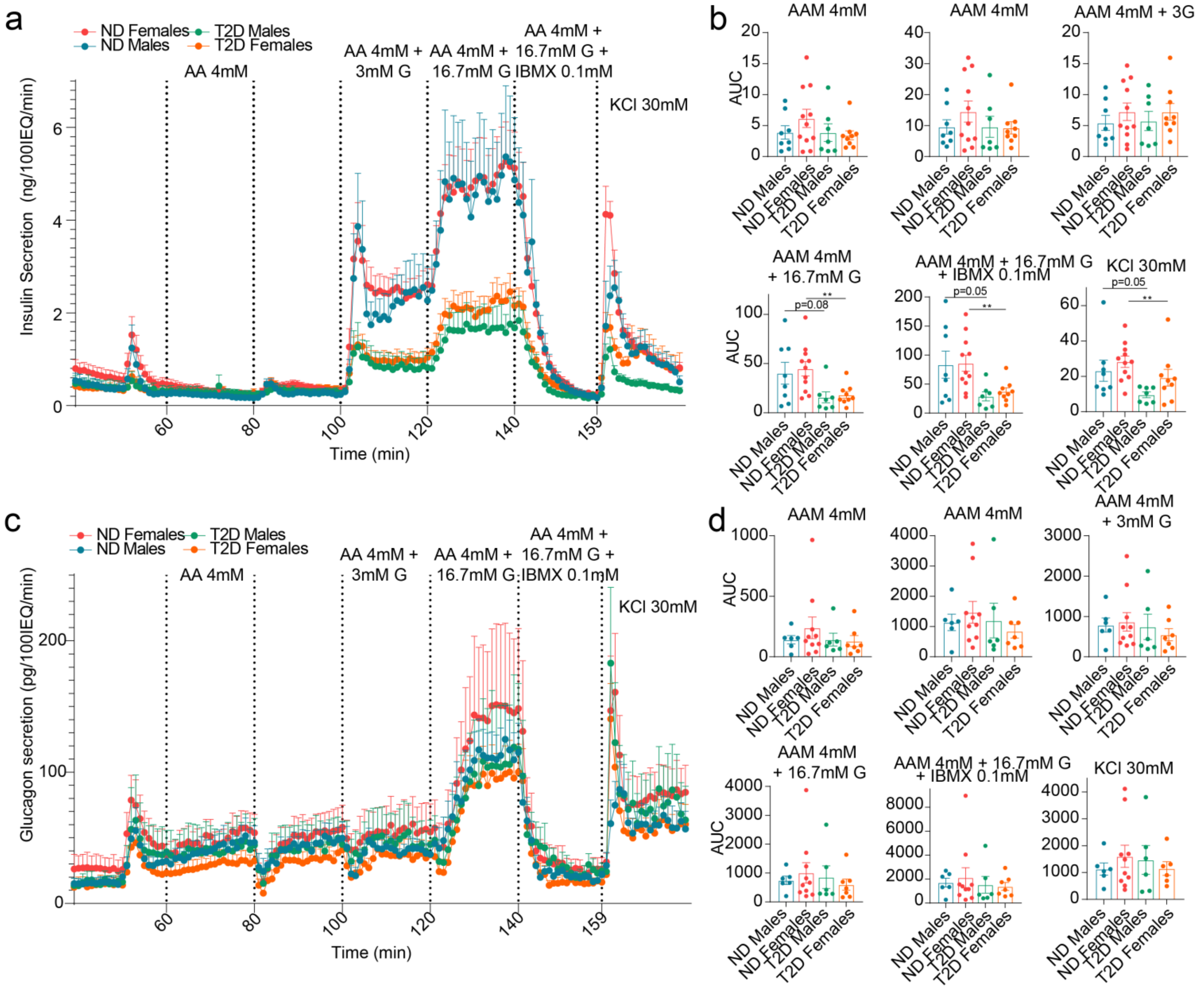


**Extended Data Fig.5 Dynamic hormone secretion profiles of HPAP type 2 diabetic donors.**

**a,** Dynamic insulin secretion assay of human islets subject to Amino acid cocktail (AA) 4mM, AA 4mM + 3mM Glucose, AA 4mM +16.7mM glucose, AA 4mM + 16.7mM Glucose + IBMX 0.1mM and KCl 30mM. Each curve represents mean of each donor type across sex and disease. **b,** Area under the curve for insulin secretion at baseline, AA 4mM, AA 4mM + 3mM Glucose, AA 4mM +16.7mM glucose, AA 4mM + 16.7mM Glucose + IBMX 0.1mM and KCl 30mM. **c,** Dynamic glucagon secretion assay of human islets subject to secretagogues as in **(a)**. **d,** Area under the curve for glucagon secretion like **(a)**. Data analyzed from HPAP dataset. n= 21 non-diabetic and n=16 T2D diabetic donors. **pvalue < 0.01

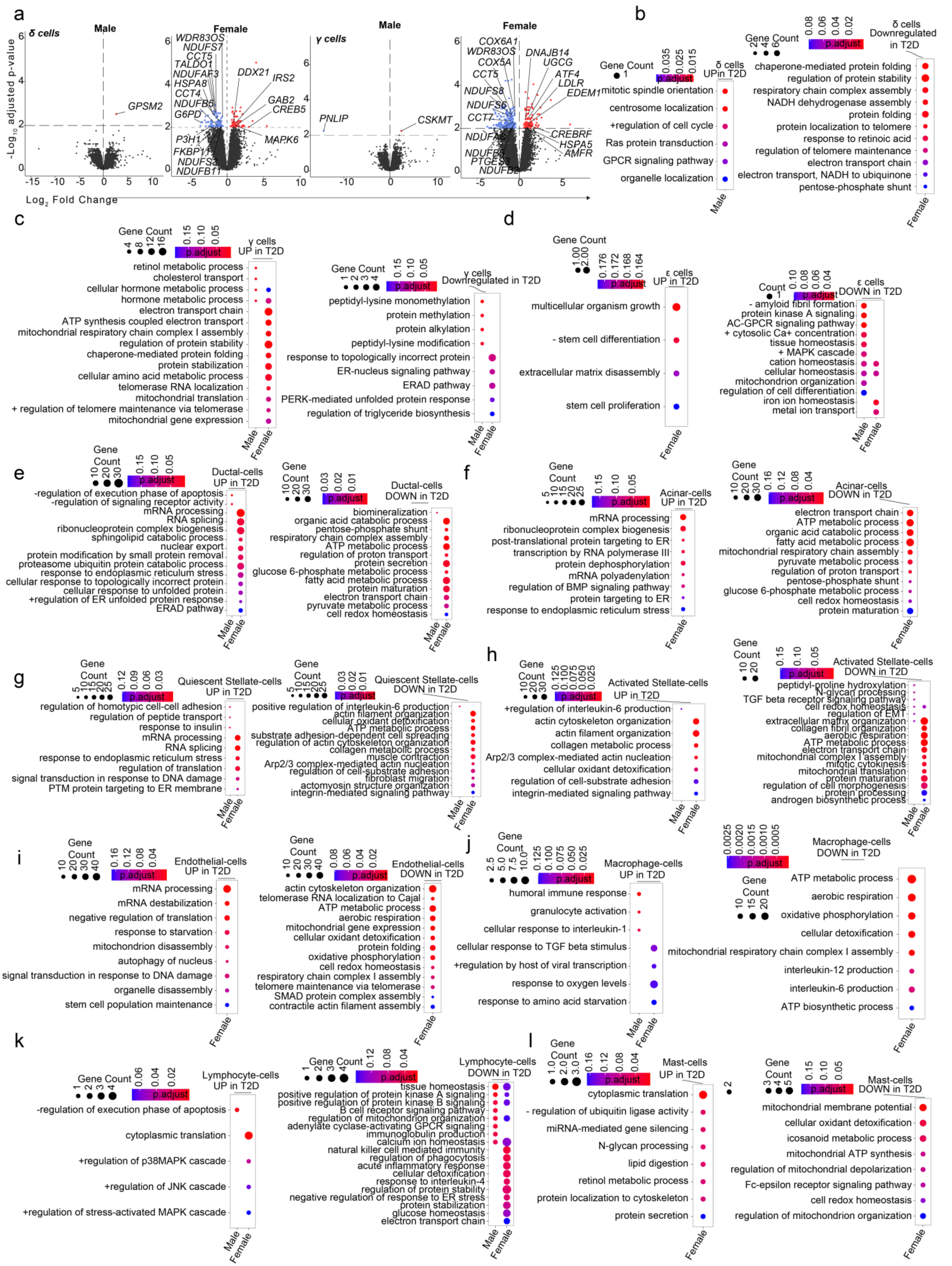

**Extended Data Fig.6 Gene ontology networks in human islet cells from type 2 diabetes donors.**

**a-h,** Gene ontology plots highlighting upregulated and downregulated pathways in T2D across sex. Pathways compare type two diabetic males to non-diabetic males and similarly for females. Data is shown for: **a,** Ductal cells; **b,** Acinar cells; **c,** Quiescent stellate cells. **d,** Activated stellate cells; **e,** Endothelial cells; **f,** Macrophage cells; **g,** Lymphocyte cells; **h,** Mast cells. DGE: FDR adjusted pval $< 0.1$ is significant, GO: FDR adjusted pval $< 0.2$ is significant, n = 36 (non-diabetic) and n = 16 T2D.
